# An inducible Cre mouse line to sparsely target nervous system cells, including Remak Schwann cells

**DOI:** 10.1101/663716

**Authors:** Darshan Sapkota, Joseph D. Dougherty

## Abstract

Nerves of the peripheral nervous system contain two classes of Schwann cells– myelinating Schwann cells that ensheath large caliber axons in myelin, and Remak Schwann cells that surround smaller axons and do not myelinate. While tools exist for genetic targeting of myelinating Schwann cells, such reagents have been challenging to generate for the Remak population in part because many of the genes that mark this population in maturity are also robustly expressed in progenitors of all Schwann cells. To circumvent this challenge, we utilized BAC transgenesis to generate a mouse line expressing a tamoxifen-inducible Cre under the control of a Remak-expressed gene promoter (*Egr1*). However, as *Egr1* is also an activity dependent gene expressed by some neurons, we flanked this Cre by flippase (Flpe) recognition target sites, and coinjected a BAC expressing Flpe under control of a pan-neuronal *Snap25* promoter, to excise the Cre transgene from these neuronal cells. Genotyping and inheritance demonstrate that the two BACs co-integrated into a single locus, facilitating maintenance of the line. Anatomical studies following a cross to a reporter line show sparse tamoxifen-dependent recombination in Remak Schwann cells within the mature sciatic nerve. However, depletion of neuronal Cre activity by Flpe is partial, with some neurons and astrocytes also showing evidence of Cre reporter activity in the central nervous system. Thus, this mouse line will be useful in mosaic loss-of-function studies, lineage tracing studies following injury, and live cell imaging studies or other experiment benefiting from sparse labeling.

## INTRODUCTION

Although glia outnumber neurons in the vertebrate nervous system and are essential for neuronal health and function, they are relatively less studied than neurons. In the peripheral nervous system (PNS), glial cells called Schwann cells are most commonly known for the formation of the myelin sheath which insulates and protects axons and ensures that nerve impulses travel quickly and efficiently^1^. Some Schwann cells however do not form the myelin sheath, and these nonmyelinating Schwann cells are among the least studied cells in the nervous system. Remak Schwann cells (RSCs), a class of nonmyelinating Schwann cells, ensheath small, 0.5–1.5 μm diameter axons, such as C fiber nociceptors in sciatic nerves and form Remak bundles^2^. It is now accepted that RSCs provide trophic support to unmyelinated axons^1^ and they are also implicated in nerve regeneration^2^ and tumorigenisis^3^. RSCs and myelinating Schwann cells share a common pool of progenitors, which express many known molecular markers of mature RSCs^4^. This has precluded the generation of RSC-specific Cre lines and thus hindered progress in understanding the function of these cells.

When a single cell type-specific *cis* regulatory element is not readily available, intersectional BAC (bacterial artificial chromosome) transgenesis can be used to moderate the expression of a transgene driven by a promiscuous *cis* regulatory element^5–7^. This is achieved by using a second transgene that precludes the expression of the first one in some cells. Most often, this is done by crossing two transgenic lines together to create a genetic gate. However, this is less efficient as only a fraction of the progeny has both transgenes. We previously demonstrated that multiple BACs can be integrated into the same locus, yet still show independent transgene expression^8^. Using a combination of these two strategies, we report a Cre/Flpe bi-transgenic mouse line that can be used to sparsely target cells in the nervous system, including RSCs in the PNS and astrocytes and neurons in the central nervous system (CNS).

## RESULTS

### Generation of *Egr1-Cre*-ER^T2^; *Snap25-Flpe* mice

To permit temporal-specificity in genetic manipulation, we utilized Cre-ER^T2^, a tamoxifen-inducible version of Cre recombinase, and drove its expression with a 92 kb BAC fragment covering early growth response 1 (*Egr1). Egr1*, an immediate early gene, is undetectable in the embryonic nervous system, and increasingly induced postnatally, with a widespread expression in the adult brain^9,10^. In the PNS, it is expressed by Schwann cell precursors during development, but confined to non-myelinating Schwann cells in the adulthood^11^. We reasoned that using a temporally specific Cre, we could avoid developmental recombination. Further, in order to restrict *Egr1*-*Cre-*ER^T2^ expression to fewer cell types, we flanked the *Cre-*ER^T2^ cassette with Flippase Recognition Targets (FRTs) and used a 61 kb BAC covering synaptosomal-associated protein 25 kDa (*Snap25*), previously shown to be neuron-specific^12^, to drive Flpe expression. Because *Snap25* is a pan-neuronal gene, we reasoned that the Cre-ER^T2^ cassette should be excised by Flpe in neurons during development. Thus, upon tamoxifen injection, Cre would only be expressed in Egr1-positive/SNAP25-negative cells (e.g. RSCs) (**Fig.1a**).

**Figure 1.**
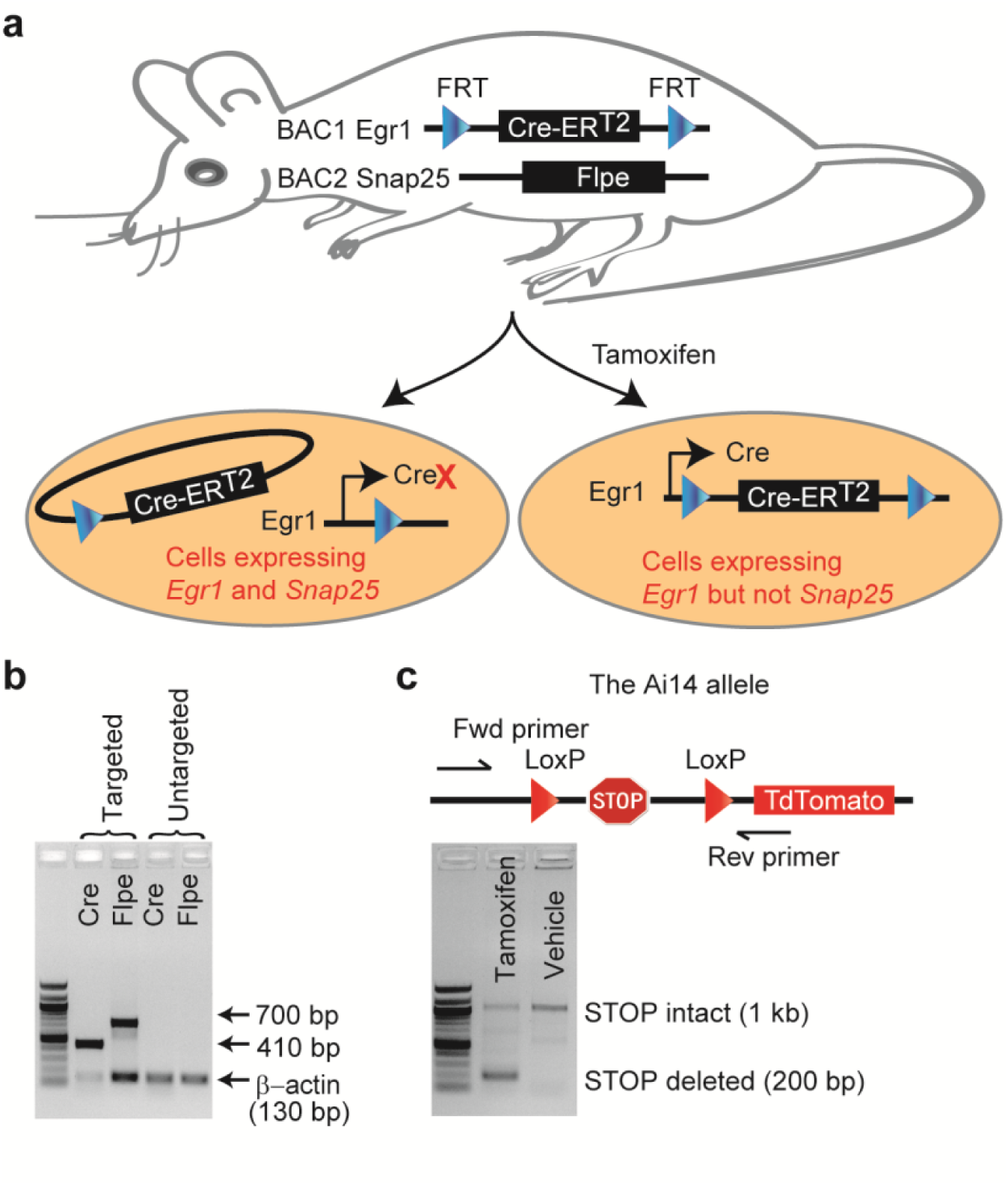
Generation of ECSF mice. **a)** Intersectional transgenesis. The two BACs shown were co-injected into mouse eggs to generate double transgenic mice. In neurons and other cells with active *Snap25* promoter, the Flpe recombinase excises the FRT-flanked Cre cassette, thus inhibiting Cre expression. Remak and other cells with active *Egr1* promoter but inactive *Snap25* promoter express Cre upon Tamoxifen treatment. **b)** PCR genotyping of founders.Representative founders with or without the Cre and Flpe transgenes are shown. **c)** Cre activity in ECSF mice. ECSF mice were crossed to Ai14 mice, which have a LoxP-flanked stop cassette excisable by Cre (upper). The ECSF; Ai14 progeny was injected with tamoxifen or vehicle, and tail preparations were PCRed using primers flanking the Ai14 LoxP sites. Excision of the stop cassette is evident in mice receiving tamoxifen but not vehicle (lower). ERT2, Estrogen Receptor; FRT, flippase recognition target.

The two BACs were co-injected into the pronuclei of fertilized mouse eggs, and the resulting progeny were PCR-genotyped for Cre and Flpe **(Fig. 1b)** to identify founders. Upon crossing the founders to wildtype C57BL/6 mice and genotyping multiple F1 mice, we found that the two transgenes were always co-inherited, suggesting that they were integrated into a single locus (data not shown). This feature avoids the need to maintain the two alleles independently and hence enhances the utility of the *Egr1-Cre*-ER^T2^; *Snap25-Flpe* (ECSF) mouse line.

We next tested if Cre is active in ECSF mice by crossing the F1 mice to Ai14 reporter mice. The Ai14 mice have a LoxP-flanked stop cassette that prevents the expression of a reporter TdTomato fluorophore unless it is excised by Cre recombinase^13^. A PCR designed to detect the excision of this stop cassette indeed showed that ECSF mice express Cre recombinase at sufficient levels to mediate recombination upon tamoxifen induction **(Fig. 1c)**.

### Identification of cells expressing Cre in ECSF mice

We first asked if RSCs show Cre activity. The ECSF;Ai14 double heterozygote mice from ECSF X Ai14 cross were injected with either tamoxifen or vehicle control (sunflower oil), and their sciatic nerves were immunostained for p75 neurotrophin receptor (p75NTR), which has been used to mark Remak bundles^14^. A robust expression of TdTomato was present in the nerves from tamoxifen-treated mice but not in those from vehicle-treated mice, evidencing a stringent inducibility of the Cre activity **(Fig. 2)**. Around 15% p75NTR-positive cells expressed TdTomato, suggesting that recombination occurred sparsely in RSCs (**Fig. 2)**.

**Figure 2.**
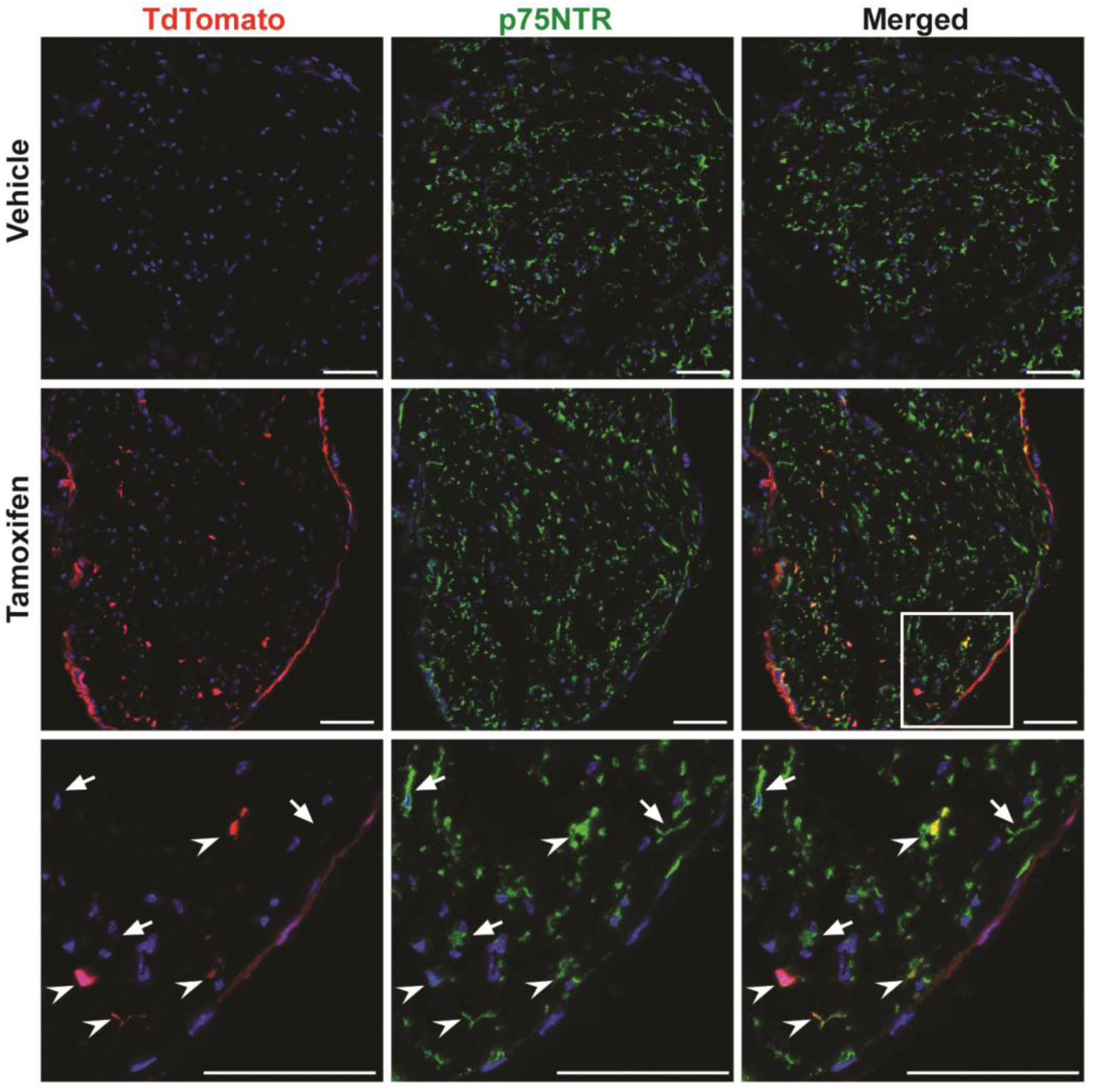
ECSF mice sparsely label Remak Schwann cells. ECSF; Ai14 mice were injected with tamoxifen or vehicle and the sciatic nerves immunostained for Remak cells (p75NTR, green) and nuclei (DAPI, blue). TdTomato is undetectable in vehicle-injected mice (top) and induced sparsely in Remak cells by tamoxifen (middle). Boxed area is enlarged to depict targeted (arrowheads) and untargeted (arrows) Remak cells (bottom). Scales bars = 50 µm.

We also stained the sciatic nerves for myelin basic protein (MBP), a marker of myelinating Schwann cells^15^. While TdTomato did not overlap with MBP, it was present in the center of ~ 10% of the myelin sheaths, suggesting Cre recombination in some axons **(Fig. 3a)**. Longitudinal sections of the nerves indeed showed a few TdTomato-expressing axons **(Fig. 3b)**. Thus, Cre activity is excluded from myelinating Schwann cells, confirming the precision of *Egr1*-*Cre*, but present sparsely in peripheral axons, suggesting an insufficiency of *Snap25*-*Flpe* activity.

**Figure 3.**
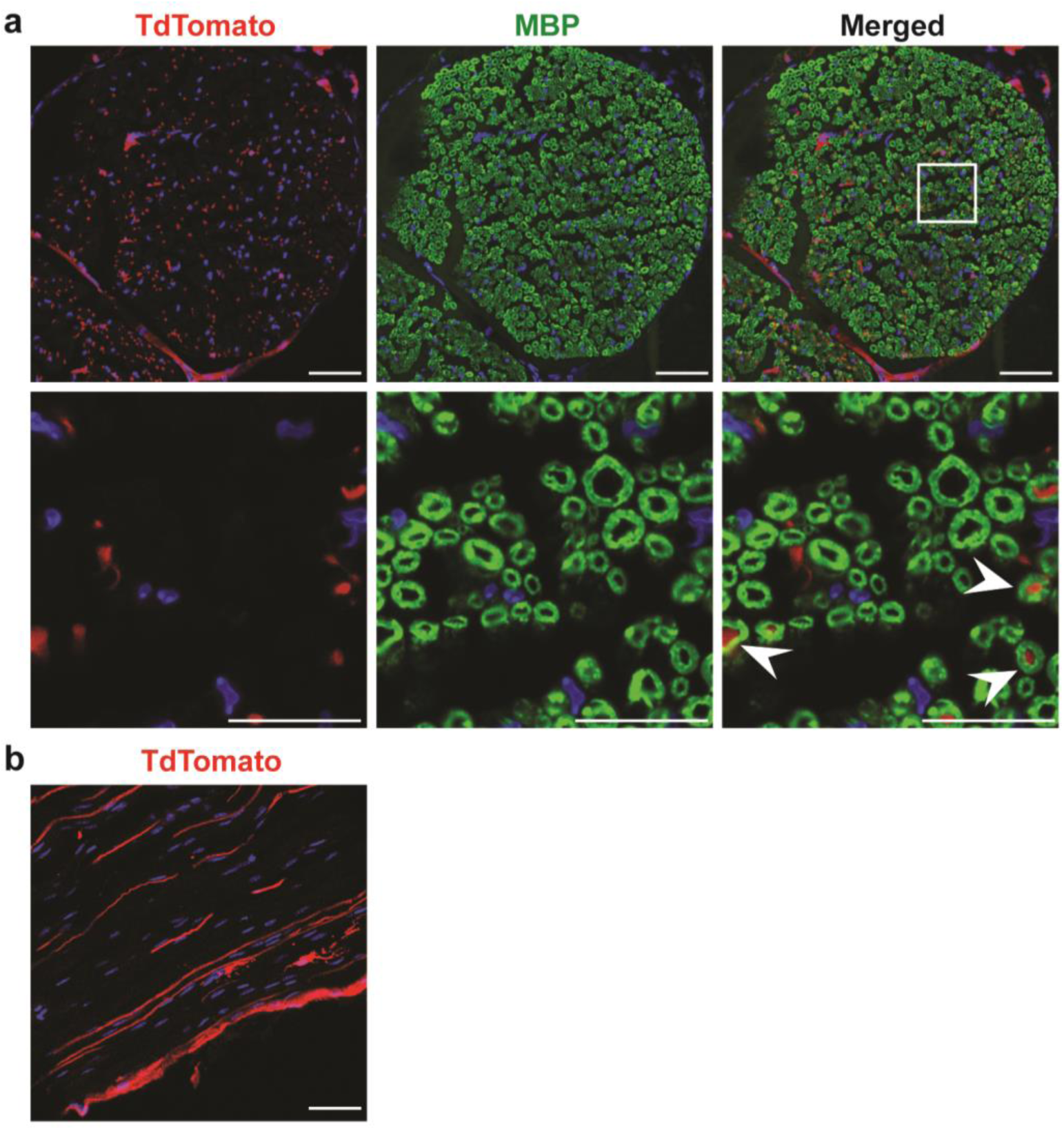
ECSF mice do not target myelinating Schwann cells. **a)** Sciatic nerves of tamoxifen-injected ECSF; Ai14 mice were immunostained for myelinating Schwann cells (MBP, green) and nuclei (DAPI, blue). Lower row are enlarged views of the boxed area. A few myelin sheaths show TdTomato-expressing axons in the center (arrowheads). **b)** Longitudinal view of the nerve shows TdTomato in axons. Scales bars = 50 µm in upper row of a and in b; 20 µm in lower row of a.

Finally we stained brain sections from tamoxifen- and vehicle-injected mice for astrocytes (Gfap) and neurons (NeuN). In Tamoxifen-injected mice, TdTomato clearly overlapped with sparse astrocytes and neurons **(Fig. 4a, b)**. Quantification in the cortex revealed 7% of Gfap positive astrocytes and 11% of neurons to be TdTomato positive. Again, TdTomato was undiscernible in the brains of vehicle-injected mice **(Fig. 4c)**.

**Figure 4.**
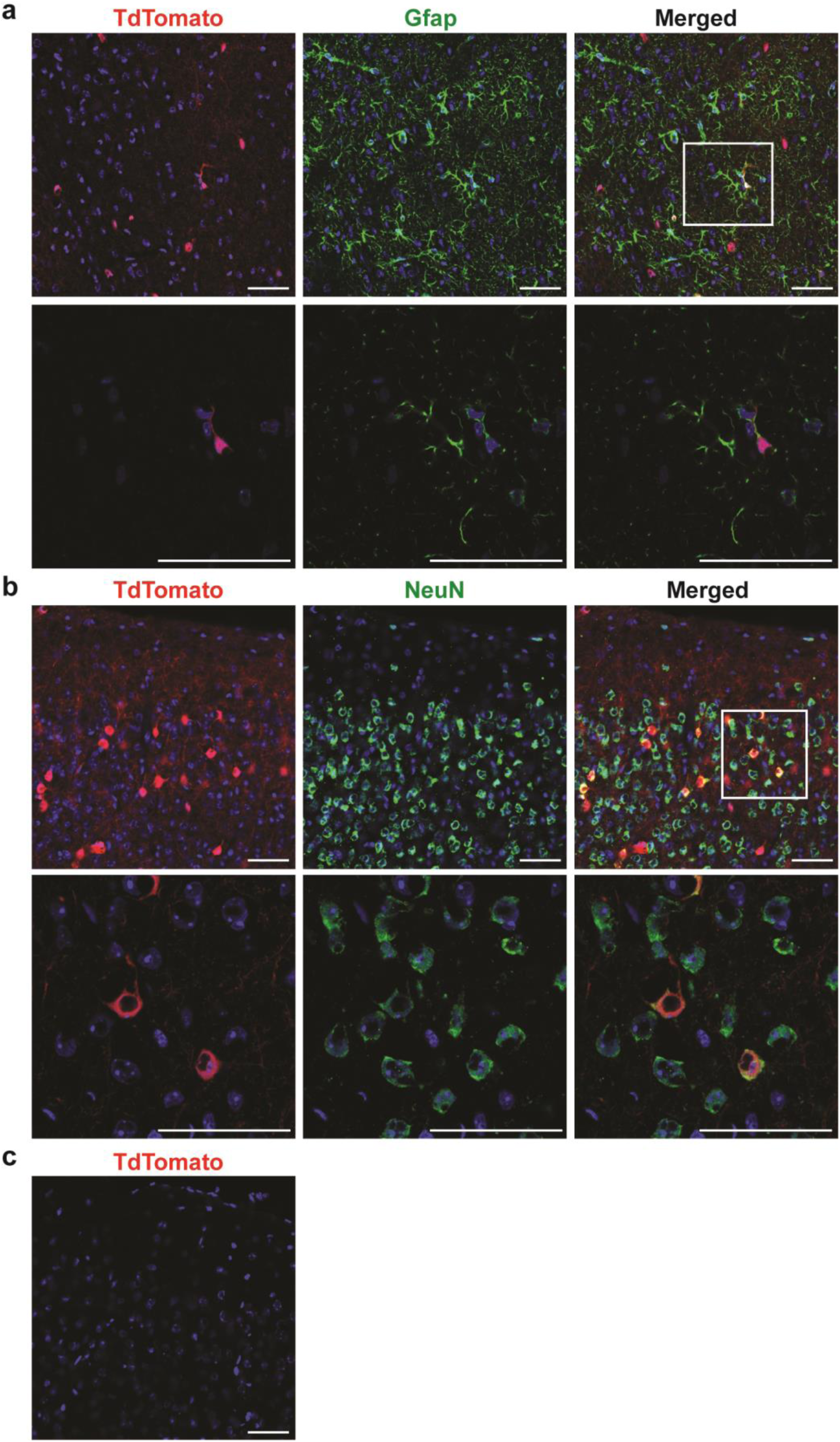
ECSF mice sparsely label cells in the brain. **a, b)** Brains from tamoxifen- and vehicle-injected ECSF; Ai14 mice were immunostained for astrocytes (Gfap, green), neurons (NeuN, green) or nuclei (DAPI, blue). Lower rows are enlarged views of the boxed regions. **c)** Cre activity is absent in the brain of vehicle-injected mice. Scales bars = 50 µm.

## DISCUSSION

We introduce the ECSF mouse to genetically target RSCs and other cell types in the nervous system. We show that cell targeting in this novel bi-transgenic mouse is sparse and stochastic. We were also able to manipulate the number of targeted cells by increasing or decreasing the number of tamoxifen injections (not shown). In principle, this mouse will be useful in loss-of-function studies when only a subset of cells needs to be studied or when possible non cell-autonomous effects arising from gene deletion need to be avoided. In addition, it will allow lineage tracing studies on RSCs, which are known to de-differentiate into repair cells following nerve injury^16^. The ECSF mouse, however, may not be ideal for studying gross pathophysiology due to a loss-of-function mutation.

Although we had expected the Cre activity to be entirely eliminated from neurons, this was not the case, possibly due to insufficient activity of the *Snap25-Flpe* BAC fragment. Yet there are occasions when a line that can reliably tag sparse neurons would be useful. For example, the Brainbow mouse has been an important reagent in studying the connectome in the brain^17^. With sparse labeling of neurons in the CNS and axons in the PNS, the ECSF mouse should be useful for similar purposes. Therefore, we have donated this mouse to the Jackson Laboratory.

## MATERIALS AND METHODS

### BAC recombination and pronuclear injection

A FRT-ER^T2^-Cre-ER^T2^-FRT cassette and an *Egr1* A box were cloned into a shuttle vector, which was then used to modify a *Egr1* BAC (clone ID: RP23-108C3) using homologous recombination as described previously^18^. Similarly, a Flpe and a *Snap25* A box were cloned into a shuttle vector, which was used to modify a Snap25 BAC (clone ID: RP23-290A18). BAC modifications were confirmed using restriction enzyme digestions followed by pulsed field gel electrophoresis. For pronuclear injections, a 92kb Egr1-FRT-ER^T2^-Cre-ER^T2^-FRT BAC fragment generated from PmeI and PacI double digestion and a 61kb Snap25-Flpe BAC fragment generated from SbfI and PmeI double digestion was purified using pulsed field gel electrophoresis followed by gel extraction (Qiagen #20021). The pronuclei of fertilized one-cell eggs of C57BL/6j mice were co-injected with a mixture of equimolar amounts of the two fragments by the Department of Pathology Knockout, Transgenic, and Microinjection Core, and then implanted into pseudopregnant foster females. The founders were genotyped using PCR. Transmission was detected in a single founder which was then crossed to C57BL/6j mice.

### Animal care

All procedures involving mice conformed to the Washington University institutional animal care and use committee. All experimental protocols were approved by the Animal Studies Committee of Washington University.

### Genotyping

Screening of ECSF founders and genotyping of their progeny were done with PCR using Cre Fwd (CCGGTCGATGCAACGAGTGATGAGGTTC), Cre Rev (GCCAGATTACGTATATCCTGG CAGCG), Flpe Fwd (CACTGATATTGTAAGTAGTTTGC) and Flpe Rev (CTAGTGCGAAGTAGTGATCAGG). Ai14 mice were PCR-genotyped as described by the Jackson Laboratory for stock number 007914. Cre recombination activity in ECSF; Ai14 mice was confirmed with PCR using the Fwd primer GCGGGCCCTAAGAAGTTCC and Rev primer TCGCCCTTGCTCACCATG to detect excision of Stop cassette.

### Tamoxifen injection

A 10 mg/ml solution of 4-hydroxy-tamoxifen (OHT) was prepared in a mixture of autoclaved sunflower oil (9 part) and ethanol (1 part) with end-to-end rotation for 1 hr at room temperature and stored at 4°C in a light-proof container. Six-week old mice were injected intraperitoneally with 2 mg (200 uL) tamoxifen for 3 subsequent days. Control mice received equal volume of the oil-ethanol vehicle. Mice were processed for immunostaining after two weeks of injections.

### Immunofluorescence staining

Mice were euthanized and perfused with 15 ml ice-cold phosphate buffer saline (PBS), followed by 20 ml 4% ice-cold paraformaldehyde in PBS. Nerves and brains were harvested; fixed in 4% ice-cold paraformaldehyde overnight; cryoprotected with 10%, 20%, and 30% ice-cold sucrose in PBS for 4 h, 4 h, and overnight, respectively; and frozen in OCT (Sakura Inc). 10 μm nerve sections and 40 μm brain sections were made for staining. Sections were blocked with 5% normal donkey serum plus 0.3% Triton® X-100 in PBS at room temperature for 1 h, incubated with primary antibody in block at 4°C overnight. Antibodies and dilutions were: rabbit anti-p75NTR (Cell Signaling Technology, 8238, 1:2000), rat anti-MBP (Biorad, MCA409S, 1:100), goat anti-Gfap (Abcam, ab53554, 1:1000), and mouse anti-NeuN (Millipore, mab377, 1:200). Following incubation with primary antibody, sections were washed three times with PBS, incubated with Alexa fluorophore-conjugated secondary antibodies (1:500, Invitrogen) in block at room temperature for 1 h, washed two times with PBS, incubated with 300nM DAPI (Sigma) at room temperature for 10 m, washed two times with PBS, and mounted for confocal imaging (Perkin Elmer). TdTomato was strong endogenously and did not require an anti-RFP antibody.

## ACKNOWLEDGEMENTS

We would like to thank Mike White for oocyte injection, Kelly Monk for helpful discussions, Michael Vasek and Claire Weichselbaum for careful editing of the manuscript, and Allison Soung for assistance with nerve dissection. This work was supported by 5R21NS083052 and 5R01NS102272.

## AUTHOR CONTRIBUTIONS

D.S. performed all experiments and wrote the paper. J.D.D. acquired the funding, supervised the project, and helped write the paper.

## REFERENCES

1. Nave, K.-A. Myelination and support of axonal integrity by glia. Nature 468, 244–252 (2010).

2. Griffin, J. W. & Thompson, W. J. Biology and pathology of nonmyelinating Schwann cells. Glia 56, 1518–1531 (2008).

3. Carroll, S. L. & Ratner, N. How Does the Schwann Cell Lineage Form Tumors in NF1? Glia 56, 1590–1605 (2008).

4. Jessen, K. R. & Mirsky, R. The origin and development of glial cells in peripheral nerves. Nat. Rev. Neurosci. 6, 671–682 (2005).

5. Dymecki, S. M., Ray, R. S. & Kim, J. C. Mapping cell fate and function using recombinase-based intersectional strategies. Methods Enzymol. 477, 183–213 (2010).

6. Taniguchi, H. et al. A Resource of Cre Driver Lines for Genetic Targeting of GABAergic Neurons in Cerebral Cortex. Neuron 71, 995–1013 (2011).

7. Plummer, N. W. et al. Expanding the power of recombinase-based labeling to uncover cellular diversity. Development 142, 4385–4393 (2015).

8. Dougherty, J. D., Zhang, J., Feng, H., Gong, S. & Heintz, N. Mouse Transgenesis in a Single Locus with Independent Regulation for Multiple Fluorophores. PLOS ONE 7, e40511 (2012).

9. Crosby, S. D. et al. Neural-specific expression, genomic structure, and chromosomal localization of the gene encoding the zinc-finger transcription factor NGFI-C. Proc. Natl. Acad. Sci. U. S. A. 89, 4739–4743 (1992).

10. Mataga, N., Fujishima, S., Condie, B. G. & Hensch, T. K. Experience-dependent plasticity of mouse visual cortex in the absence of the neuronal activity-dependent marker egr1/zif268. J. Neurosci. Off. J. Soc. Neurosci. 21, 9724–9732 (2001).

11. Topilko, P. et al. Differential regulation of the zinc finger genes Krox-20 and Krox-24 (Egr-1) suggests antagonistic roles in Schwann cells. J. Neurosci. Res. 50, 702–712 (1997).

12. Dougherty, J. D. et al. Candidate pathways for promoting differentiation or quiescence of oligodendrocyte progenitor-like cells in glioma. Cancer Res. 72, 4856–4868 (2012).

13. Madisen, L. et al. A robust and high-throughput Cre reporting and characterization system for the whole mouse brain. Nat. Neurosci. 13, 133–140 (2010).

14. Michellin, L. B. et al. Leprosy patients: neurotrophic factors and axonal markers in skin lesions. Arq. Neuropsiquiatr. 70, 281–286 (2012).

15. Salzer, J. L. Schwann Cell Myelination. Cold Spring Harb. Perspect. Biol. a020529 (2015). doi:10.1101/cshperspect.a020529

16. Jessen, K. R. & Mirsky, R. The repair Schwann cell and its function in regenerating nerves. J. Physiol. 594, 3521–3531 (2016).

17. Livet, J. et al. Transgenic strategies for combinatorial expression of fluorescent proteins in the nervous system. Nature 450, 56–62 (2007).

18. Gong, S., Yang, X. W., Li, C. & Heintz, N. Highly efficient modification of bacterial artificial chromosomes (BACs) using novel shuttle vectors containing the R6Kgamma origin of replication. Genome Res. 12, 1992–1998 (2002).

